# The Volume of Mitochondria Inherited Impacts mtDNA Homeostasis in Budding Yeast

**DOI:** 10.1101/2025.03.25.645216

**Authors:** Michael W. Ray, WeiTing Chen, Chengzhe Duan, Guadalupe Bravo, Kyle Krueger, Erica M. Rosario, Alexis A. Jacob, Laura L. Lackner

## Abstract

Most eukaryotic cells maintain mitochondria in well-distributed, reticular networks. The size of the mitochondrial network and copy number of its genome scale with cell size. However, while the size scaling features of mitochondria and their genome are interrelated, the fitness consequences of this interdependence are not well understood. We exploit the asymmetric cell division of budding yeast to test the hypothesis that mitochondrial scaling with cell size impacts mitochondrial DNA (mtDNA) function. We find that the volume of mitochondria inherited by daughter cells affects the ability of cells to maintain functional mtDNA; daughter cells that inherit a significantly reduced volume of mitochondria have an increased frequency of losing respiratory competence. In cells with such mitochondrial inheritance defects, mtDNA integrity can be maintained by upregulating mtDNA copy number. Collectively, these data support a bet-hedging model whereby the faithful inheritance of an adequate volume of mitochondria ensures enough mtDNA copies are transmitted to daughter cells to counteract pre-existing and/or inevitable mtDNA mutations.

**Summary:** Ray et al. demonstrate that the volume of mitochondria inherited impacts mtDNA homeostasis in the model system budding yeast. They propose a model by which inheritance of an adequate mitochondrial volume results in the transmission of sufficient mtDNA copies to counteract existing and/or inevitable mutations.

## INTRODUCTION

Decades of work have resolved the strategies eukaryotic cells use to shape and position their defining, essential organelle, the mitochondrion. Mitochondria in most eukaryotic cells exist in dynamic, reticular networks that sit well-distributed throughout the cell. However, some cell types adopt distinct mitochondrial morphologies for unique purposes, such as the discrete mitochondrial structures used by neurons for axonal transport (Pekkurnaz and Wang, 2022). The overall morphology of mitochondria is determined by the conserved mechanisms of mitochondrial dynamics (division/fusion), anchoring, and trafficking (Lackner, 2014; Friedman and Nunnari, 2014). These activities shape and position mitochondria as well as impact the function of the organelle, highlighting the intimate connections between mitochondrial morphology and function.

Mitochondria cannot form *de novo*. Thus, mitochondria must be partitioned during cell division—a process known as mitochondrial inheritance—to ensure cell survival. Mitochondrial dynamics, anchoring, and trafficking have all been shown to contribute to mitochondrial inheritance (Bockler et al., 2017; Higuchi-Sanabria et al., 2016; Lackner et al., 2013). In addition, mitochondrial biogenesis—which requires the addition of lipids, proteins, and nucleic acids to existing compartments—is critical to ensure mitochondrial content is not progressively lost following each successive round of cell division. Thus, dividing cells must regulate mitochondrial content throughout the cell cycle. A seminal study in budding yeast demonstrated that mitochondrial volume scales with cell size (Rafelski et al., 2012), and similar cell size scaling of mitochondria has been observed in higher-order eukaryotes (Miettinen and Bjorklund, 2016). However, how cells sense and regulate mitochondrial volume in accordance with their cell size remains unclear. This is an especially complicated challenge for mitochondria, which contain an outer mitochondrial membrane (OMM), a cristae-forming inner mitochondrial membrane (IMM), and a matrix that houses mtDNA. mtDNA is packaged into proteinaceous structures called nucleoids (Miyakawa, 2017), and the number of nucleoids and copies of mtDNA have been shown to scale with mitochondrial length and cell size, respectively, in yeast (Jajoo et al., 2016; Osman et al., 2015; Seel et al., 2023). The molecular mechanisms that coordinate mitochondrial size, mtDNA copy number, and cell size remain largely unknown.

In asymmetrically dividing cells, such as budding yeast, mitochondria must be actively transported to the daughter cell prior to cytokinesis to ensure faithful mitochondrial inheritance. Mmr1 and Ypt11 are partially redundant adaptors for the type V myosin motor, Myo2, that facilitates the actin-dependent transport of mitochondria from the mother cell to the growing bud (Boldogh et al., 2004; Chernyakov et al., 2013; Eves et al., 2012; Itoh et al., 2004; Itoh et al., 2002). Mmr1 can directly bind the outer mitochondrial membrane (Chen et al., 2018) and Myo2 (Itoh et al., 2004; Eves et al., 2012), bridging the interaction between the organelle and motor protein. The mechanism of Ypt11-mediated mitochondrial inheritance, however, is unclear. Ypt11 is a Rab GTPase that binds the cargo-binding domain (CBD) of Myo2 at a site distinct from Mmr1 (Eves et al., 2012; Tang et al., 2019). The GTPase activity of Ypt11 facilitates the trafficking of mitochondria from mother to bud (Lewandowska et al., 2013) but the molecular basis for its mitochondrial association is not known. Ypt11 also functions in Golgi and ER inheritance (Arai et al., 2008; Swayne et al., 2011), and whether its role in these processes is related to its role in mitochondrial inheritance is unclear. Deletion of either adaptor alone results in a mitochondrial inheritance delay (Boldogh et al., 2004; Chen et al., 2018; Chernyakov et al., 2013; Itoh et al., 2004; Itoh et al., 2002) while the simultaneous depletion of both adaptors is lethal, as mitochondrial inheritance is inhibited (Chernyakov et al., 2013; Kraft and Lackner, 2017). Thus, the presence of at least one of these adaptors is required for mitochondrial inheritance and, consequently, cell viability.

Given that mitochondrial inheritance is an essential prerequisite to deliver a template upon which the mitochondrial biogenesis machinery can propagate the compartment for proper mitochondria to cell size scaling, we reasoned that mitochondrial inheritance would also impact mtDNA scaling and function. Here, we exploit the asymmetric nature of budding yeast cell division to test this hypothesis. We first quantified the contributions of Mmr1 and Ypt11 to the volume of mitochondria inherited. We then exploited the quantified differences in mitochondrial inheritance to demonstrate that the volume of mitochondria inherited affects the ability of cells to maintain functional mtDNA in a gradated manner; when daughter cells inherit a reduced volume of mitochondria, cells have an increased frequency of losing respiratory competence. Our data support a bet-hedging model in which the inheritance of an adequate amount of mitochondria by newborn daughter cells enables the transmission of enough functional mtDNA copies to counteract pre-existing and/or inevitable critical mutations.

## RESULTS AND DISCUSSION

### Mitochondrial inheritance adaptors contribute unequally to the inheritance of mitochondrial volume by the daughter cell

Previous studies have demonstrated that cells lacking either Mmr1 or Ypt11 display a delay in mitochondrial inheritance during budding (Boldogh et al., 2004; Chen et al., 2018; Chernyakov et al., 2013; Itoh et al., 2004; Itoh et al., 2002). We reproduced these findings using the standard assay in the field in which cells are binned into small/large bud categories and scored for the presence of mitochondria in the bud (Fig. 1 A). However, a caveat of this scoring method is that the amount of mitochondria a bud inherits is not considered in the quantification. Indeed, clear qualitative differences in mitochondrial volume that were not captured by this method of scoring were readily observed in large buds of Δ*mmr1* and Δ*ypt11* cells (Fig. 1 B). Therefore, to take the amount of mitochondria into account, we adapted a pipeline to rigorously assess the volume of mitochondria in buds over the progression of budding. Using live-cell 3D confocal microscopy of mitochondrial matrix-targeted DsRed (mito-dsRED) and MitoGraph, a published tool for accurately measuring mitochondrial volume in a region of interest (ROI) (Rafelski et al., 2012; Viana et al., 2015), we plotted the volume of mitochondria in the buds of wild type (WT), Δ*mmr1*, and Δ*ypt11* cells against the bud-to-mother size (b:m) ratio, a measure of cell cycle progression (Fig. 1 C).

**Figure 1.**
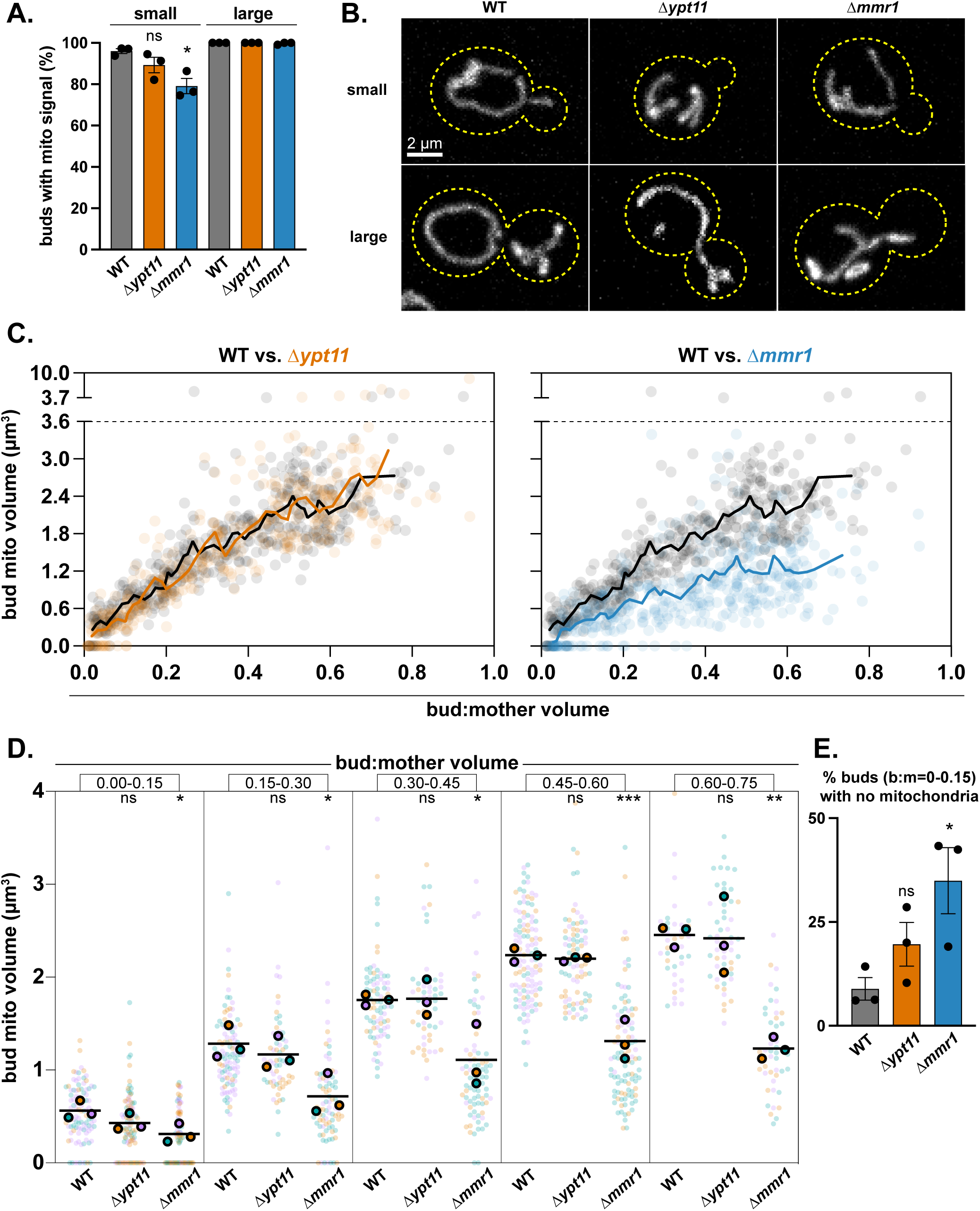
Cells lacking Mmr1 have a more severe mitochondrial inheritance defect than cells lacking Ypt11. Wild-type (WT), Δ*ypt11*, and Δ*mmr1* cells expressing mito-dsRED were grown in SCD at 24°C and analyzed by fluorescence microscopy. The same dataset was used for the entire figure. Only budded cells were analyzed. (A) Quantification of percent buds of a given category (cut-off for small vs. large is a b:m volume ratio of 1:3) with mitochondrial signal present. Each black dot represents the mean for one (of 3 total) biological replicates with >35 cells analyzed per replicate in a given category. Colored bars show the grand mean with error bars representing ± SEM. p-values are in comparison to WT of the small bud category. (B) Representative images for each category in A shown as whole-cell maximum projections with the cell wall outlined with a yellow dashed line. Bar, 2 µm. (C) Quantification of mitochondrial volume in the bud (µm^3^) plotted against the b:m volume ratio. WT (gray) vs. Δ*ypt11* (orange) on the left. WT (gray) vs. Δ*mmr1* (blue) on the right. Pooled data from 3 biological replicates (>80 cells per replicate, >350 cells total per genotype) shown as one data point per cell. Lines show 15-point moving averages. Y-axes are broken in two for efficient visualization of a minimal number of real but high-value data points. (D) Data from C was collapsed into 0.15-range bins for b:m volume ratio values up to 0.75 (chosen as upper limit to ensure data analysis with sufficient n values). Bins within b:m volume ratio values of 0-0.6 contain ≥15 data points per replicate, and the 0.6-0.75 bin contains ≥7 data points per replicate. The Y-axis is limited to 4 µm^3^ (compare with C) for visualization purposes. Individual biological replicates are shown in different colors. Individual biological replicate means are shown in black-outlined circles of the given color, and the grand mean is shown as a black line. p-values are in comparison to WT of the same bin. (E) Quantification of percent buds in the smallest b:m volume bin (b:m = 0-0.15) from D that lack mitochondrial signal. Each black dot represents the mean for one (of 3 total) biological replicates. Colored bars show grand mean with error bars representing ± SEM. p-values are in comparison to WT (* p < 0.05; ** p < 0.01; **** p < 0.0001; ordinary one-way ANOVA multiple comparisons).

Consistent with Rafelski et al., 2012, the volume of mitochondria in the buds of WT cells scaled positively with the b:m ratio. The distribution of bud mitochondrial volume for Δ*ypt11* cells appeared remarkably similar to that of WT cells, with exception to the two earliest b:m bins (b:m = 0.00-0.15 and 0.15-0.30), in which Δ*ypt11* cells exhibited a decrease in bud mitochondrial volume (Fig. 1, C and D). Furthermore, Δ*ypt11* cells showed an increase in % buds that lack mitochondria in comparison to WT cells for the earliest b:m bin (Fig. 1E). Thus, the reduction in bud mitochondrial volume in small buds of Δ*ypt11* cells was fully restored as budding progressed (Fig. 1, C and D), consistent with the results of Rafelski et al. 2012. In striking contrast, the bud mitochondrial volume of Δ*mmr1* cells was significantly lower than that of both WT and Δ*ypt11* cells across all stages of budding (Fig. 1, C and D). In addition, the number of buds in the earliest b:m bin that lacked mitochondria was increased in Δ*mmr1* cells compared to both WT and Δ*ypt11* cells (Fig.1, E).

To determine if the differences in bud mitochondrial volume were due to differences in bud size, bud mitochondrial volume was normalized to the volume of the bud (referred to as “normalized bud mitochondrial volume”) (Fig. S1 A). Comparisons of this measure between WT and either Δ*ypt11* or Δ*mmr1* cells were similar to comparisons of raw bud mitochondrial volume in Fig. 1 C, indicating that the differences in bud mitochondrial volume could not be attributed to differences in bud volume. These results demonstrate that cells lacking Mmr1 have a more severe mitochondrial inheritance defect than cells lacking Ypt11. While the subtle defect in mitochondrial inheritance in Δ*ypt11* cells is restored as budding progresses, cells lacking Mmr1 have a significantly lower mean bud mitochondrial volume than both WT and Δ*ypt11* cells that persists throughout budding.

One obvious question these data raise is whether the mitochondrial inheritance defects in the Mmr1 and Ypt11 mutants lead to an accumulation of mitochondria in the mother. To address this, we repeated the mitochondrial volume assay using ROIs that capture the mother of each mother-bud pair. We found that the mother mitochondrial volume normalized to mother volume (referred to as “normalized mother mitochondrial volume”) for WT, Δ*mmr1*, and Δ*ypt11* cells were largely maintained over bud progression relative to themselves and each other (Fig. S1 B). Thus, despite the observed differences in normalized bud mitochondrial volume across these strains, their normalized mother mitochondrial volumes were similar over the course of the cell cycle. Together our data indicate that Mmr1 and Ypt11 make unequal contributions to mitochondrial inheritance; Mmr1 contributes to the inheritance of a greater mitochondrial volume during the cell cycle than Ypt11.

### Severe disruptions in mitochondrial inheritance result in post-cytokinetic mitochondrial scaling defects in daughter cells

In WT cells, mitochondrial volume scales with cell size (Rafelski et al., 2012). To test how this relationship is affected when mitochondrial inheritance is disrupted, we exploited our robust dataset from the mitochondrial inheritance quantification pipeline shown in Fig. 1. We first plotted the total mitochondrial volume normalized to cell volume for budded cells, which represents an unsynchronized global population (Fig. 2 A). The mean of this measure of normalized total mitochondrial volume for Δ*mmr1* cells was subtly lower in comparison to that of WT and Δ*ypt11* cells, suggesting that severe mitochondrial inheritance defects can cause small perturbations to the known mitochondria-to-cell size scaling relationship.

**Figure 2.**
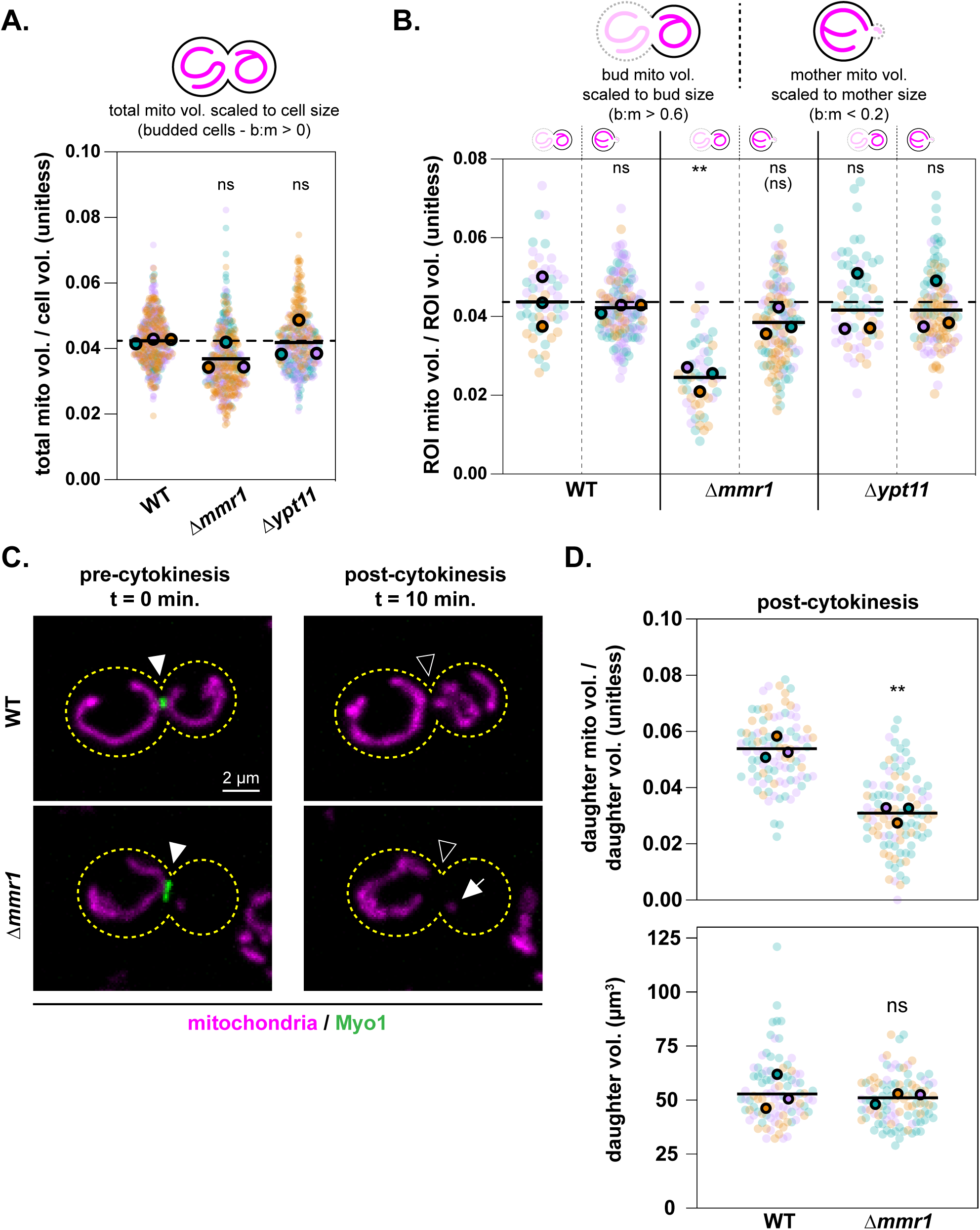
Daughter cells lacking Mmr1 undergo cytokinesis with a lower normalized bud mitochondrial volume. (A-B) Using the dataset from Fig. 1 C, mitochondrial volume (µm^3^) was normalized to cell volume (µm^3^), bud volume (µm^3^), or mother volume (µm^3^), creating a unitless scaling ratio. Individual biological replicates (3 total) are shown in different colors. Individual biological replicate means are shown in black-outlined circles of the given color, and the grand mean is shown as a black line. The grand mean of the associated WT value is shown as a horizontal dashed line in black. Panel A shows the quantification of normalized whole-cell mitochondrial volume using budded cells only (b:m > 0) as a representation of a vegetative population. Each replicate contains >80 cells. p-values are in comparison to WT (* p < 0.05; ** p < 0.01; **** p < 0.0001; ordinary one-way ANOVA multiple comparisons). For each genotype shown in Panel B, normalized mitochondrial volume ratios of two different regions of interest (ROIs) are shown on either side of the vertical dashed line: normalized bud mitochondrial volume for late-stage budded cells where the b:m > 0.6 (left) and normalized mother mitochondrial volume for early-stage budded cells where the b:m < 0.2 (right). >10 cells were analyzed for each replicate of the late-stage bud measure (>50 cells total). >20 cells were analyzed for each replicate of the early-stage mother measure (>110 cells total). p-values are in comparison to normalized bud mitochondrial volume (b:m > 0.6) of WT cells, and p-value in parenthesis compares both measures of Δ*mmr1* cells (* p < 0.05; ** p < 0.01; **** p < 0.0001; ordinary one-way ANOVA multiple comparisons). (C-D) WT and Δ*mmr1* cells expressing mito-dsRED and Myo1-GFP were imaged every 10 min for 3 h at room temperature in SCD-containing imaging dishes to identify and analyze cytokinetic events. Representative images of mitochondria and Myo1 in pre- and post-cytokinetic cells are shown in C as whole-cell maximum projections. Cell walls are outlined with dashed yellow lines. Filled arrowheads point at accumulated Myo1 signal on the mother-bud neck at the last timepoint before such signal is lost (“pre-cytokinesis”). Empty arrowheads point to the mother-bud neck void of Myo1 signal at the first timepoint in which Myo1 signal is not present at this location (“post-cytokinesis”). The arrow points to the minimal amount of mitochondria in the daughter cell of the representative Δ*mmr1* image post-cytokinesis. Bar, 2 µm. For at least 17 cells per biological replicate (>95 cells total from 3 biological replicates that are shown in different colors), the first timepoint in which Myo1-GFP signal was lost from the mother-bud neck was analyzed and called “post-cytokinesis”. This provided a measure of both normalized mitochondrial volume in the daughter cell (D, upper panel) and daughter volume (D, lower panel) following cytokinesis. Individual biological replicate means are shown in black-outlined circles of a given color and the grand mean is shown as a black line. p-values are in comparison to WT (* p < 0.05; ** p < 0.01; **** p < 0.0001; two-tailed unpaired *t* test).

To better understand where in the cell cycle the mitochondria-to-cell size scaling relationship is affected in our adaptor mutants, we next compared mitochondrial volume normalized to size for specific ROIs for WT, Δ*mmr1*, and Δ*ypt11* cells at specific stages of budding: late-stage budded cells (b:m > 0.6) and early-stage budded cells (b:m < 0.2). These stages represent populations about to undergo and having recently undergone cytokinesis, respectively. For late-stage budded cells, the normalized bud mitochondrial volume for WT and Δ*ypt11* cells was indistinguishable while Δ*mmr1* cells exhibited a significant decrease in normalized bud mitochondrial volume (Fig. 2 B). Interestingly, while the normalized mother mitochondrial volume of early-stage budded Δ*mmr1* cells was also reduced in comparison to WT, the difference between WT and Δ*mmr1* cells at this stage of the cell cycle was smaller in magnitude. Given that late-stage buds turn into new mothers, and new mothers comprise at least 50% of the cell population at any given time in an unsynchronized population of cells, these data suggest that Δ*mmr1* cells undergo cytokinesis with a significantly lower normalized bud mitochondrial volume and that mitochondria-to-cell size scaling is partially restored in the newborn cell by the time a bud emerges.

To more definitively demonstrate that Δ*mmr1* cells undergo cytokinesis with a reduced normalized bud mitochondrial volume, we turned to live-cell 4D confocal microscopy. We used strains expressing endogenous Myo1 fused to yEGFP as a marker to visually track cytokinesis since the protein accumulates at the incipient bud site and disappears rapidly upon cytokinesis (Li et al., 2021; Lippincott and Li, 1998). We tracked the cell wall, mitochondria, and Myo1 in WT and Δ*mmr1* cells every 10 minutes and identified instances in which Myo1 disappeared from the mother-bud neck. Then, using the first timepoint that lacked an accumulated Myo1 signal, we quantified mitochondrial and daughter cell volume and verified that Δ*mmr1* cells indeed undergo cytokinesis with a reduced normalized daughter mitochondrial volume compared to WT (Fig. 2, C and D). Importantly, the size of Δ*mmr1* buds at the time of cytokinesis did not differ from WT, indicating a true defect in the raw volume of mitochondria inherited (Fig. 2 D). Therefore, cells with defected mitochondrial inheritance can undergo cytokinesis with a minimal amount of mitochondria in the bud, consistent with previous observations that mitochondrial inheritance is not required for cytokinesis (Chernyakov et al., 2013). Despite their mitochondrial inheritance defect, our data suggest that Δ*mmr1* daughter cells are able to partially restore their normalized mitochondrial volume by the time they form a new bud (Fig. 2 B). Notably, the growth rate of Δ*mmr1* cells was similar to WT (Fig. S2 A), suggesting this mitochondrial inheritance defect does not significantly alter cell cycle progression. Together, these findings suggest homeostatic mechanisms exist for cells to not only scale the amount of mitochondria with cell size as previously reported (Rafelski et al., 2012) but also correct perturbations in this scaling after cytokinesis.

### Mitochondrial inheritance adaptors contribute unequally to the maintenance of mtDNA integrity

Both mitochondrial volume and mtDNA copy number scale with cell size, and nucleoid number scales with mitochondrial network length (Osman et al., 2015; Rafelski et al., 2012; Seel et al., 2023). Thus, leveraging the differences in mitochondrial inheritance and the resulting impact on mitochondria-to-cell size scaling we observed between Δ*mmr1* and Δ*ypt11* cells, we next examined the effect of altered mitochondrial inheritance on mtDNA maintenance. To assess the ability of Δ*mmr1* and Δ*ypt11* cells to maintain functional mtDNA, we measured their petite frequencies (Fig. 3 A). This assay distinguishes respiratory competent (*rho^+^*) from respiratory incompetent (petite) cells and is a standard method to assess mtDNA maintenance (Ogur and St John, 1956; Shadel, 1999). Strikingly, we found that, after 14 hours of fermentative growth (which does not require functional mtDNA and, therefore, allows for the propagation of mtDNA mutations), Δ*mmr1* cells have an increased petite frequency (∼45%) compared to WT (∼10%) and Δ*ypt11* cells (∼10%). Notably, similar to WT and Δ*ypt11* cells, 100% of Δ*mmr1* cells grown in fermentation conditions contained DAPI-marked mtDNA signal (Fig. 3 B). Therefore, the respiratory incompetence of petite Δ*mmr1* cells was likely due to mtDNA mutation (*rho^-^* cells) and not mtDNA loss (*rho^0^* cells). Together, these results indicate that Mmr1 contributes to the maintenance of functional mtDNA, a process for which Ypt11 is dispensable. Furthermore, these findings highlight the differential contributions made by the partially redundant mitochondrial inheritance adaptors to cellular homeostasis.

**Figure 3.**
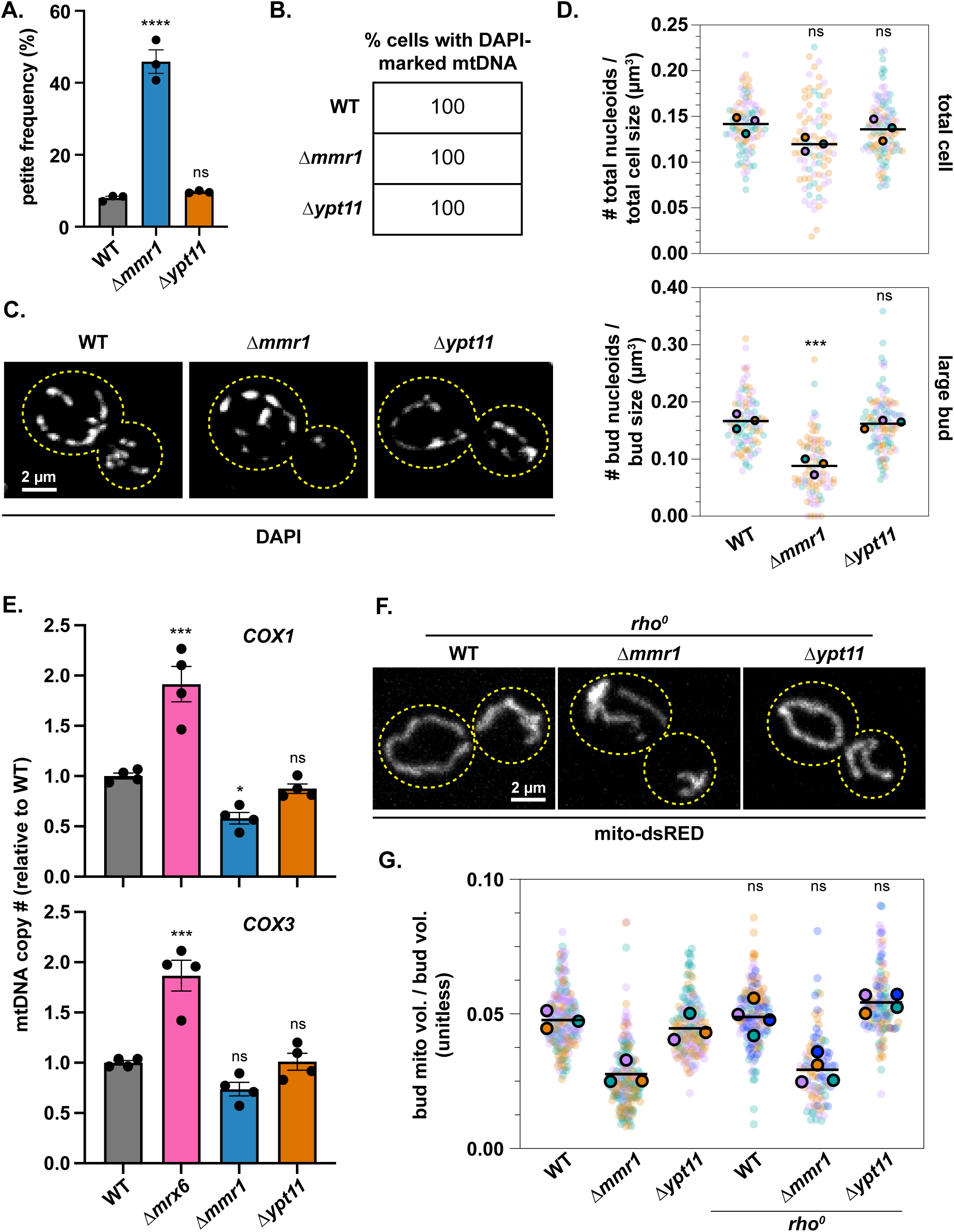
Cells lacking Mmr1 display a defect in the maintenance of mtDNA integrity. (A) Petite frequency (%) of wild-type (WT), Δ*mmr1*, and Δ*ypt11* cells after 14 h in fermentation (YPD) conditions in which respiration-compromising mtDNA mutations can propagate and petite cells can survive. Each black dot represents an individual biological replicate (3 total) in which >100 colonies were counted. Colored bars show grand mean with error bars representing ± SEM. p-values are in comparison to WT (* p < 0.05; ** p < 0.01; **** p < 0.0001; ordinary one-way ANOVA multiple comparisons). (B-D) WT, Δ*mmr1*, and Δ*ypt11* cells were grown in SCD at 30°C, stained with DAPI, and analyzed by fluorescence microscopy. Panel B shows the quantification of the percent of cells that contain DAPI-marked mtDNA (and are therefore either *rho^+^* or *rho^-^*). Representative images of DAPI-stained cells are shown in C as whole-cell maximum projections. Cell walls are shown as yellow dashed lines. Bar, 2 µm. Panel D shows the quantification of the number of DAPI-marked nucleoids normalized to a given region of interests’ volumes: either total cell (b:m > 0.4 as a proxy for a vegetative population, upper panel) or large buds (b:m > 0.4, lower panel). Individual biological replicates are shown in different colors. Individual biological replicate means (3 total) are shown in black-outlined circles of the given color, and the grand mean is shown as a black line. Each replicate contains at least 15 cells (>85 cells total). p-values are in comparison to WT (* p < 0.05; ** p < 0.01; **** p < 0.0001; ordinary one-way ANOVA multiple comparisons). (E) qPCR analysis of mtDNA alleles (*COX1* and *COX3*) in WT, Δ*mmr1*, Δ*ypt11*, and Δ*mrx6* cells. mtDNA copy number relative to WT is shown. 2-3 technical replicates per biological replicate (4 biological replicates total per genotype, shown individually as black dots) were included. Colored bars show grand mean with error bars representing ± SEM. p-values are in comparison to WT (* p < 0.05; ** p < 0.01; **** p < 0.0001; ordinary one-way ANOVA multiple comparisons). (F) Representative images of mitochondria from a *rho^0^* background for the given genotypes are shown as whole-cell maximum projections. Cell walls are shown as yellow dashed lines. Bar, 2 µm. (G) Normalized mitochondrial volume of large buds (b:m > 0.4) of *rho^0^* cells expressing mito-dsRED grown in SCD at 24°C and analyzed by fluorescence microscopy. WT/Δ*mmr1*/Δ*ypt11* data are from the same dataset shown in Fig. 1. Individual biological replicates (3 total for WT/Δ*mmr1*/Δ*ypt11* and 4 total for all other genotypes) are shown in different colors. Individual biological replicate means are shown in black-outlined circles of a given color, and the grand mean is shown as a black line. Each biological replicate contains at least 18 cells (>95 cells total). p-values compare a given genotype to its *rho^0^* counterpart (* p < 0.05; ** p < 0.01; **** p < 0.0001; ordinary one-way ANOVA multiple comparisons).

### The distribution and scaling of mtDNA relative to mitochondria are retained in cells with severe mitochondrial inheritance defects

We next sought to determine whether the defect in the maintenance of mtDNA integrity in Δ*mmr1* cells could be attributed to differences in nucleoid number and distribution. To this end, we stained cells with DAPI and used live-cell 3D confocal microscopy to visualize and measure the number of nucleoids in WT, Δ*mmr1*, and Δ*ypt11* cells (Fig. 3, C and D). DAPI-marked nucleoids appeared as well-distributed punctate structures in Δ*mmr1* and Δ*ypt11* cells, phenocopying those of WT cells. We found that the number of nucleoids per cell normalized to cell volume was decreased slightly in Δ*mmr1* cells (85% of the WT value based on the grand mean of 3 biological replicates; Fig. 3 D, upper panel) and correlated with the slight decrease in normalized total mitochondrial volume in Δ*mmr1* cells (88% of the WT value based on the grand mean of 3 biological replicates; Fig. 2 A). Furthermore, the difference in the number of bud nucleoids normalized to bud volume in Δ*mmr1* cells with a b:m > 0.4 (53% of the WT value based on the grand mean of 3 biological replicates; Fig. 3 D, lower panel) correlated with the difference in normalized bud mitochondrial volume that was observed in Δ*mmr1* cells of the same b:m bin (58% of the WT value based on the grand mean of 3 biological replicates; Fig. 3 G, left 2 columns). For this and subsequent analyses of buds, we focused on cells with a b:m > 0.4 since this stage more accurately reflects the amount of mitochondria daughter cells inherit upon cytokinesis.

Given the polyploid nature of mtDNA and its relationship to the amount of mitochondria (Osman et al., 2015; Seel et al., 2023), we also quantified mtDNA copy number in our adaptor mutants. We performed qPCR on two mtDNA alleles that sit at spatially distinct loci, *COX1* and *COX3,* in WT, Δ*mmr1*, and Δ*ypt11* cells as well as in Δ*mrx6* cells, which exhibit an increase in mtDNA copy number (Goke et al., 2020), as a positive control. Similar to the comparisons of normalized total mitochondrial volume between the strains, the relative mtDNA copy number of Δ*mmr1* cells was reduced in comparison to WT and Δ*ypt11* cells (Fig. 3 E). In summary, our combined imaging and qPCR analyses demonstrate that known scaling features of mtDNA are preserved in Δ*mmr1* and Δ*ypt11* cells. Namely, in these strains, nucleoids sit well-distributed as discrete puncta and mtDNA copy number is correlated with the amount of mitochondria. Therefore, the defect in the maintenance of mtDNA integrity in cells lacking Mmr1 is likely not due to perturbations in nucleoid distribution or the mtDNA-mitochondria scaling relationship.

### The presence of mtDNA does not affect mitochondrial inheritance

Given that Δ*mmr1* cells have an increased frequency of becoming petite, we wanted to determine if respiratory incompetence contributed to the mitochondrial inheritance defect observed in these cells. We repeated our volume-based mitochondrial inheritance assay in WT, Δ*mmr1*, and Δ*ypt11* cells that lack mtDNA (*rho^0^*). We found that the normalized bud mitochondrial volume for each genotype examined was not significantly altered in the absence of mtDNA (Fig. 3, F and G). These results indicate that the presence of mtDNA does not affect the amount of mitochondria inherited and suggest that mitochondrial inheritance is not impacted by the respiratory competence of the cell in fermentative conditions. Consequently, the increased petite frequency of Δ*mmr1* cells does not appear to contribute to their mitochondrial inheritance defect.

### The volume of mitochondria inherited correlates with the maintenance of mtDNA integrity

To further interrogate the role of Mmr1 in the maintenance of mtDNA integrity, we examined the petite frequency of cells expressing Mmr1 mutants that disrupt the interaction between Mmr1 and mitochondria (*mmr1*Δ*mito-BD* and *mmr1^4E^*) (Fig. 4 A; mutants fully described in Table S1) (Chen et al., 2018). We found that the petite frequency of these Mmr1 mutants phenocopied that of cells lacking Mmr1. These results suggest that the role of Mmr1 in the maintenance of mtDNA integrity depends on the ability of Mmr1 to interact with mitochondria.

**Figure 4.**
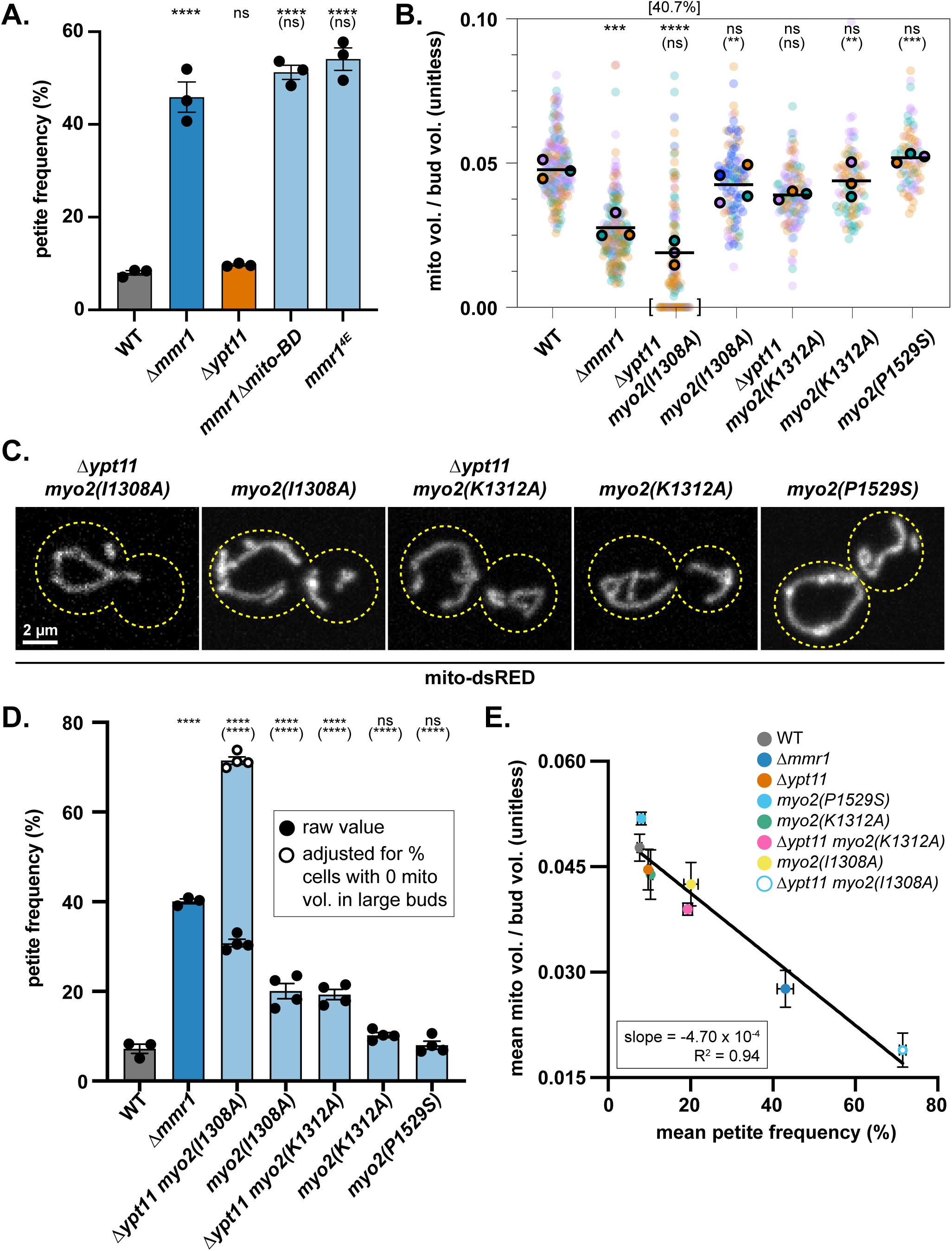
Titrating the volume of mitochondria inherited to determine the relationship between the volume of mitochondria inherited and the maintenance of mtDNA integrity. (A) Petite frequency (%) of *mmr1* mutants with disrupted mitochondria interaction after 14 h in fermentation (YPD) conditions, in which respiration-compromising mtDNA mutations can propagate and petite cells can survive. The petite frequency of wild-type (WT), Δ*mmr1*, and Δ*ypt11* cells is duplicated from Fig. 3 A for ease of comparison. Each black dot represents an individual biological replicate (3 total) in which >100 colonies were counted. Colored bars show grand mean with error bars representing ± SEM. p-values are in comparison to WT, and p-values in parenthesis are in comparison to Δ*mmr1* (* p < 0.05; ** p < 0.01; **** p < 0.0001; ordinary one-way ANOVA multiple comparisons). (B) Quantification of normalized mitochondrial volume in large buds (as in Fig. 3 G) of *myo2* single point mutant strains expressing mito-dsRED. WT and Δ*mmr1* data are reproduced from Fig. 3 G for ease of comparison. Individual biological replicates [3 total except for *myo2(I1308A)* for which there are 4 to maintain a comparable n] are shown in different colors. Individual biological replicate means are shown in black-outlined circles of the given color and the grand mean is shown as a black line. The data points sitting at a value of 0 as well as the percent of cells with the value of 0 for the pooled data are indicated by the brackets at the bottom and top of the graph, respectively. Each biological replicate contains ≥19 cells (≥75 cells total). p-values are in comparison to WT, and p-values in parenthesis are in comparison to Δ*mmr1* (* p < 0.05; ** p < 0.01; **** p < 0.0001; ordinary one-way ANOVA multiple comparisons). (C) Representative fluorescence microscopy images for the cells quantified in 4B. Cell walls are outlined with yellow dashed lines. Bar, 2 µm. (D) Petite frequency (%) of *myo2* single point mutant strains determined and represented as in 4A. 4 biological replicates are represented for the *myo2* point mutant strains. The open circles indicate adjusted petite frequency values: the percent of large buds with 0 mitochondrial volume in B was added to the petite frequency (%) to account for petite cells that die. p-values are in comparison to WT, and p-values in parenthesis are in comparison to Δ*mmr1* (* p < 0.05; ** p < 0.01; **** p < 0.0001; ordinary one-way ANOVA multiple comparisons). (E) Grand mean values of the normalized mitochondrial volume of large buds from B and Fig. 3 G are plotted against the mean petite frequency (%) values from Fig. 3 A and Fig. 4 D with error bars representing ± SEM. Slope and R^2^ are derived from linear regression for which the line of best fit is shown in black. Open circle represents data using experimentally adjusted values as described in 4D.

Given that both the Mmr1-mitochondria and Mmr1-Myo2 interaction are required for Mmr1-mediated mitochondrial transport (Chen et al., 2018), we wanted to determine if perturbing mitochondrial transport using mutations that disrupt the Mmr1-Myo2 interaction would also impact the maintenance of mtDNA integrity. We turned to three previously characterized point mutants in the Myo2-CBD that specifically disrupt the Mmr1-Myo2 interaction to varying degrees— *myo2(P1529S)*, *myo2(K1312A),* and *myo2(I1308A)* (Eves et al., 2012). The purpose of using these Myo2 mutants was two-fold. First, they allowed us to disrupt Mmr1-mediated mitochondrial transport without perturbing the Mmr1 protein to examine whether the mtDNA integrity defect could be ascribed to a function of Mmr1 outside of transport. Second, we reasoned that these mutants would provide varied mitochondrial inheritance defects, enabling us to test the idea that the volume of mitochondria inherited during the cell cycle correlates with the maintenance of functional mtDNA.

First, we examined the growth of cells expressing these Myo2 mutants from the endogenous *MYO2* locus in the presence and absence of *YPT11* and found that only *myo2(I1308A)* Δ*ypt11* cells exhibited a severe growth defect (Fig. S2 A). Next, we investigated mitochondrial inheritance in cells expressing the Myo2 point mutants from the endogenous locus in the presence of Ypt11 for *myo2(P1529S)* and in both the presence and absence of Ypt11 for *myo2(I1308A)* and *myo2(K1312A)* (Fig. 4, B and C). We quantified the normalized bud mitochondrial volume in cells with a b:m > 0.4. *myo2(I1308A)* cells exhibited a mitochondrial inheritance defect that was further exacerbated in the absence of Ypt11. Indeed, a high percentage of *myo2(I1308A)* Δ*ypt11* cells lacked any mitochondrial signal in the bud, consistent with the growth defect observed for this strain. Similarly, the subtle mitochondrial inheritance defect in cells expressing *myo2(K1312A)* was exacerbated in the absence of Ypt11. Cells expressing *myo2(P1529S)* did not show a defect in mitochondrial inheritance. Therefore, we successfully generated strains that exhibit a range of normalized large bud mitochondrial volumes that span from mean values lower than Δ*mmr1* cells to mean values similar to WT and Δ*ypt11* cells. Importantly, this suite of mutants maintained mtDNA copy numbers at a level close to or above Δ*mmr1* cells, suggesting that these mutants do not result in unexpected mtDNA loss (Fig. S2 B).

We next examined the petite frequency of these strains as a measure of the maintenance of mtDNA integrity (Fig. 4 D). A complicating factor for this experiment is that any cells that do not inherit mitochondria will die and not be included in the quantification despite being petite by definition, which is relevant for *myo2(I1308A)* Δ*ypt11* cells. Therefore, for this strain, we plotted both the experimentally determined petite frequencies as well as petite frequencies adjusted for the fact that ∼40% of large buds lack mitochondria (Fig. 4 B; see brackets) and, therefore, will likely produce daughter cells that lack mitochondria and, consequently, die. Consistent with the mitochondrial inheritance data, we found that *myo2(I1308A)* cells have a petite frequency between that of WT and Δ*mmr1* cells while the petite frequency of *myo2(P1529S)* and *myo2(K1312A)* cells approached WT levels. Consistent with our observed defects in mitochondrial inheritance, the petite frequencies of the *myo2(K1312A)* and *myo2(I1308A)* point mutants increased in the absence of Ypt11. *myo2(I1308A)* Δ*ypt11* cells exhibited the highest petite frequency of the mutants tested (Fig. 4 D), with an adjusted mean value above that of Δ*mmr1* cells, and the lowest normalized bud mitochondrial volume of the mutants tested (Fig. 4 B), with a mean below that of Δ*mmr1* cells.

Together, these data indicated that not only the Mmr1-mitochondria interaction but also the Mmr1-Myo2 interaction are involved in the role of Mmr1 in the maintenance of mtDNA integrity; thus, Mmr1-mediated mitochondrial transport appears to be integral in the protein’s contribution to this process. Furthermore, these data suggested an inverse correlation between petite frequency and the volume of mitochondria inherited. To better visualize this relationship, we plotted the grand mean value of the normalized bud mitochondrial volume against the mean petite frequency in all strains examined and found a striking inverse linear correlation (R^2^=0.94, Fig. 4 E). These data suggest that the ability of cells to maintain functional mtDNA over generations is impacted by the volume of mitochondria inherited in a gradated manner.

### Increasing mtDNA copy number reduces the loss of respiratory competence in cells lacking Mmr1

Given that nucleoids sit evenly spaced within the mitochondrial matrix, the number of mtDNA copies a daughter cell receives is tied to the volume of mitochondria inherited (Roussou et al., 2024). Therefore, we sought to determine whether the impact of mitochondrial volume inherited on the maintenance of mtDNA integrity is related to the amount of mtDNA inherited.

To address this, we asked whether increasing mtDNA copy number in cells with defective mitochondrial inheritance would reduce the rate at which the cells lose respiratory competence. We turned to three mutants that increase mtDNA copy number; these include deletions of *SML1*, which encodes a ribonucleotide reductase inhibitor (Taylor et al., 2005), *MRX6*, which encodes a putative regulator of mtDNA replication (Goke et al., 2020), and *CIM1*, which encodes a mitochondrial HMG-box protein (Schrott and Osman, 2023). As expected, we found that the deletion of each gene in Δ*mmr1* cells increased mtDNA copy number in comparison to Δ*mmr1* alone (Fig. 5 A). Furthermore, deleting these genes in Δ*mmr1* cells partially rescued the petite frequency of Δ*mmr1* cells in a statistically significant manner (Fig. 5 B). This rescue was not due to an increase in the volume of mitochondria inherited, since the normalized bud mitochondrial volume for large buds of all three double mutants phenocopied that of Δ*mmr1* (Fig. 5, C and D). To confirm that these double mutants transmit the expected number of nucleoids to buds compared to Δ*mmr1* alone, we again imaged DAPI-stained live cells to measure the number of nucleoids in large buds normalized to bud size (Fig. 5, E and F). Similar to the mitochondrial inheritance data, we found that the number of nucleoids in large buds normalized to the bud volume of the double mutants phenocopied that of Δ*mmr1* cells. This is consistent with the uncoupling of nucleoid number and mtDNA copy number previously reported for Δ*mrx6* and Δ*cim1* cells; while mtDNA copy number is increased in these cells, nucleoid number does not increase, indicating an increase in mtDNA copy number per nucleoid (Goke et al., 2020; Schrott and Osman, 2023). Thus, our data are consistent with the idea that in comparison to Δ*mmr1* cells, the number of nucleoids inherited does not change in the double mutants, but the number of mtDNA copies inherited is increased. Taken together, these data suggest that the mechanism by which the inheritance of mitochondrial volume impacts the maintenance of mtDNA integrity relates to the number of copies of mtDNA transmitted to daughter cells during cell division.

**Figure 5.**
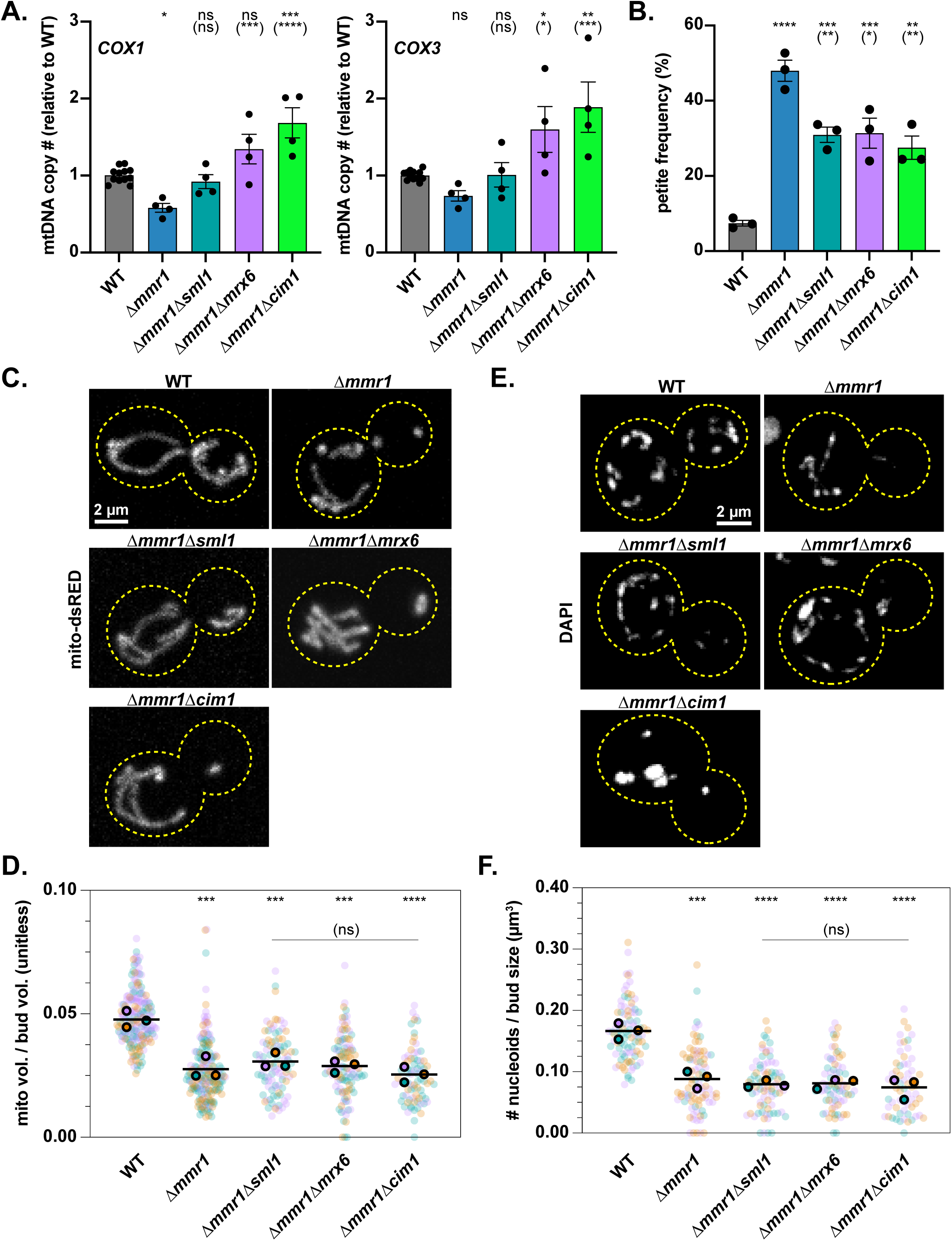
Increasing mtDNA copy number suppresses the petite frequency of Δ*mmr1* cells. (A) Cells of the indicated genotype were subjected to qPCR analysis of mtDNA alleles as in Fig. 3 E. WT and Δ*mmr1* data from Fig. 3 E are reproduced in these graphs for ease of comparison. Individual biological replicates are shown as dots. Colored bars show grand mean with error bars representing ± SEM. p-values are in comparison to WT. p-values in parenthesis are in comparison to Δ*mmr1* (* p < 0.05; ** p < 0.01; **** p < 0.0001; ordinary one-way ANOVA multiple comparisons). (B) Petite frequency (%) was determined and represented as in Fig. 3 A. p-values are in comparison to WT and p-values in parenthesis are in comparison to Δ*mmr1* (* p < 0.05; ** p < 0.01; **** p < 0.0001; ordinary one-way ANOVA multiple comparisons). (C) Representative images of mito-dsRED in indicated strains. Whole-cell maximum intensity projections are shown. Cell walls are outlined with yellow dashed lines. Bar, 2 µm. (D) Quantification of normalized mitochondrial volume in large buds (as in Fig. 3 G) of the indicated strains. WT and Δ*mmr1* data are reproduced from Fig. 3 G for ease of comparison. Each biological replicate contains at least 20 cells (>75 cells total). p-values are in comparison to WT. p-values in parenthesis are in comparison to Δ*mmr1*. (* p < 0.05; ** p < 0.01; **** p < 0.0001; ordinary one-way ANOVA multiple comparisons). (E) Representative images of DAPI-marked nucleoids in the indicated strains. Whole-cell maximum intensity projections are shown. Cell walls are outlined with yellow dashed lines. Bar, 2 µm. (F) Quantification of bud nucleoid # normalized to bud volume in large buds of indicated strains was performed and is represented as in Fig. 3 D (lower panel). WT and Δ*mmr1* data are reproduced from Fig. 3 D for ease of comparison. Each biological replicate contains ≥ 13 cells (≥ 62 cells total). p-values are in comparison to WT. p-values in parenthesis are in comparison to Δ*mmr1*. (* p < 0.05; ** p < 0.01; **** p < 0.0001; ordinary one-way ANOVA multiple comparisons).

### The inheritance of an adequate mitochondrial volume enables the transmission of sufficient mtDNA copies to maintain mtDNA function

By titrating the volume of mitochondria inherited using various mutations in the mitochondrial inheritance pathway, we demonstrated that the amount of mitochondria inherited impacts the maintenance of functional mtDNA in a gradated manner (Fig. 4 E). Mitochondria occupy ∼5% of the volume of WT buds about to undergo cytokinesis (Fig. 4 B), and for every ∼1% loss in the volume of the bud that is occupied by mitochondria, the frequency of losing respiratory competence increases by 20% for cells grown in the conditions tested (Fig. 4 E).

We reason that inheriting a nominal volume of mitochondria and, therefore, fewer copies of mtDNA increases the burden of maintaining functional gene expression from the transmitted copies of mtDNA. Cells have been shown to rely upon a threshold mtDNA copy number of functional mtDNA to maintain respiration and mtDNA function itself (Filograna et al., 2021; Smith et al., 2024). The even distribution of mtDNA-containing nucleoids across a mitochondrial network therefore likely necessitates the partitioning of adequate mitochondrial volumes over cell cycles to meet this threshold. The idea that the volume of mitochondria inherited affects mtDNA function was reported in a recent computational study that suggested reduced inheritance of mitochondrial volume could drive mtDNA homoplasmy (Roussou et al., 2024). Such take-over of one mtDNA form, if critically mutated, could contribute to the loss of respiratory function we see in cells with defective mitochondrial inheritance. Our data support a bet-hedging model whereby the inheritance of an adequate volume of mitochondria ensures enough functional mtDNA copies are transmitted to counteract existing and/or inevitable detrimental mtDNA mutations. A similar bet-hedging model for mtDNA distribution was reported in the distant fungi, *Schizosaccharomyces pombe* (Jajoo et al., 2016). mtDNA replication is highly error-prone (Anderson et al., 2020). mtDNA copy number, therefore, becomes important to ameliorate defects associated with heteroplasmic mtDNA mutations and perhaps to provide access to functional mtDNA molecules for homology-directed repair (Filograna et al., 2019; Filograna et al., 2021; Jiang et al., 2017; Nishiyama et al., 2010; Wisniewski et al., 2024; Zwonitzer et al., 2024). Indeed, we found that increasing mtDNA copy number with three separate gene deletions suppressed the petite frequency of Δ*mmr1* cells without rescuing their mitochondrial inheritance defect (Fig. 5). Given the conserved features of mtDNA distribution, future work will uncover whether mitochondrial volume partitioning during the cell cycle impacts mtDNA function in other eukaryotes. Furthermore, uncovering the mechanism(s) by which cells co-regulate their mitochondrial scaling with cell size and mtDNA copy number provides ample ground for future study.

## MATERIALS AND METHODS

### Strains and Plasmids

All strains and primers used in this study are cataloged in Supplemental Tables 1 and 2, respectively. All plasmids can be found in Supplemental Table 3. W303 (*ade2–1; leu2–3; his3– 11, 15; trp1–1; ura3–1; can1–100*) was used as the wild-type background for all strains used (Thomas and Rothstein, 1989). All new strains were constructed via either PCR-based targeted homologous recombination and/or mating followed by sporulation and tetrad analysis. Primer sets used to amplify deletion cassettes for generating knock-out strains and to amplify C-terminal gene-tagging cassettes were generated using common sequences published for such purposes (Goldstein and McCusker, 1999; Janke et al., 2004; Longtine et al., 1998; Sheff and Thorn, 2004; Sikorski and Hieter, 1989). Myo2 point mutants were generated by first knocking out one allele of *MYO2* in a W303 diploid (given that *MYO2* is essential) followed by i) homologous recombination-based insertion of PCR products amplified from plasmids containing the Myo2 single point mutations and selection cassette into the genome at the Δ*myo2* allele (Eves et al., 2012) and then ii) sporulation and tetrad dissection. Isolates were verified to contain the correct mutation by sequencing. To generate *rho^0^* cells (Fox et al., 1991), cells were grown in yeast extract/peptone with 2% (wt/vol) dextrose (YPD) plus 25 µg/ml ethidium bromide for 24 h at 30°C and were used to inoculate a second culture in the same medium. Following another 24 h of growth, the culture was streaked on YPD plates. Single colonies were isolated and tested for lack of respiratory growth on yeast extract/peptone with 3% (vol/vol) ethanol and 3% (vol/vol) glycerol (YPEG) media. All strains used for imaging mitochondria were transformed with pLL806 digested with Bsp119I. All strains and plasmids are available upon request to the corresponding author.

### Petite frequency

Petite frequency assays were performed using the established colorimetric distinction between respiration-competent (*rho^+^*) and respiration-incompetent (petite) cells which, as incipient cells, form red and white colonies, respectively, when grown on fermentative plates with low adenine (Shadel, 1999). This assay relies upon the *ade2* mutation, which our W303 strain background contains. Petite frequency was defined as the percent of all colonies that were scored that were petite.

Cells were grown on YPEG plates followed by overnight growth in liquid YPEG at 24°C or 30°C. Next, overnight cultures were diluted into fresh liquid YPEG and grown to log-phase at 24°C or 30°C. These log-phase cultures were further diluted into fresh liquid YPD to maintain log growth for 14 h at 30°C, corresponding to ∼11 doublings. Cells were then diluted before being plated and grown on YPD without supplemented adenine at 30°C before being scored for petite frequency.

### Quantitative PCR

Real-time quantitative PCR (qPCR) was performed on two alleles, *COX1* and *COX3*, sitting at distinct loci on the mitochondrial chromosome to account for possible site-specific mutations, such as deletions, in our tested strains. Strains used for qPCR contained no genetic markers or imaging-based genetic fusions beyond those used for knock-out cassettes to avoid potential artifacts. Cells were grown to log-phase in SCD [synthetic complete medium plus 2% (wt/vol) dextrose with 2x adenine at pH 6.4]. Total DNA from 5 mL of log-phase cells was isolated using the MasterPure^TM^ Yeast DNA Purification Kit from BioSearch Technologies. DNA concentration quantitation was performed on a DeNovix DS-11+ Spectrophotometer. Samples were diluted in sterile double-distilled water to 100 ng/mL for qPCR analysis. Applied Biosystems^®^ PowerUp^TM^ SYBR^TM^ green master mix (REF# A25742) along with primers indicated in Supplementary Table 2 were used to amplify *COX1* (mtDNA), *COX3* (mtDNA), and *ACT1* (as a normalizer for nuclear DNA). Samples were run in either technical triplicate or duplicate on MicroAmp^®^ optical 96-well reaction plates with optical adhesive covers (Applied Biosystems^®^ REF#s, respectively: N8010560, 4360954) on an Applied Biosystems^®^ QuantStudio 3 thermocycler. Each plate run contained two WT biological replicates for normalization; as such, WT mtDNA levels were quantified with a greater number of biological replicates than experimental strains, for which 3-4 biological replicates were measured. The ΔΔCt method was used to quantify mtDNA levels relative to wild-type based on the measured threshold Ct value from which 2^-ΔΔCt^ was calculated using the following formula: ΔΔCt = ΔCt_test strain – ΔCt_control average of wild-type (where ΔCt_test strain = Ct_COX1/3 of a given biological replicate – Ct_ACT1 of the same biological replicate; and where ΔCt_control average = average of two biological replicate measures of ΔCt_COX1/3 of a WT biological replicate – ΔCt_ACT1 of the same WT biological replicate) (Livak and Schmittgen, 2001).

### Imaging

For mitochondrial volume analysis, cells were grown to log-phase in SCD at 24°C then incubated at room temperature on optical imaging dishes from MatTek (Part No. P35G-0.170-14-C) treated with 2 mg/mL Concanavalin A to adhere cells. Dishes were washed with SCD before adding additional SCD for live-cell imaging. All images were acquired on a Nikon Spinning Disk Confocal System fitted with a CSU-W1 dual-spinning disk unit (Yokogawa) using a 60X (NA-1.42) objective and a Hamamatsu ORCA Fusion Digital CMOS camera. 3D z-stacks with a step size of 0.2 µm for both brightfield and relevant fluorescent channel(s) were acquired. Nikon Elements software was used to capture images, which were adjusted linearly for brightness and contrast as well as cropped in FIJI (Schindelin et al., 2012). Images were assembled in Illustrator (Adobe Systems). Non-deconvolved images are shown and used for all analyses.

For post-cytokinetic mitochondrial volume analysis using Myo1-GFP, cells were imaged every 10 min for 3 h to capture multiple cytokinetic events.

For DAPI analysis, log-phase cultures grown in SCD at 24°C were incubated with DAPI at a final concentration of 2.5 µg/mL for 30 minutes at room temperature prior to washing with SCD, resuspending in SCD, then incubating cells on optical imaging dishes from MatTek (Part No. P35G-0.170-14-C) treated with 2 mg/mL Concanavalin A to adhere cells. Dishes were then washed with SCD before adding additional SCD for live-cell imaging.

### Image quantification

For mitochondrial volume analysis, images were prepared for MitoGraph in FIJI. The ellipse tool was used to draw ROIs around mothers and buds using the brightfield channel. The non-deconvolved channel containing the z-stack of mito-dsRED signal was max-projected manually or using the *GenMaxProjs.ijm* plug-in. Then, individual ROI files were generated using the *CropCells.ijm* plug-in. Finally, MitoGraph v3.0 (https://github.com/vianamp/MitoGraph) was run on Macintosh Terminal using the parameters in the following code-line: ./MitoGraph -xy 0.111 -z 0.2 (Viana et al., 2015).

For DAPI analysis, the ellipse tool in FIJI was used as for the mitochondrial volume analysis above to isolate mother and bud ROIs. The “Find Maxima” function was used on maximum projections of the DAPI images for these individual ROIs. Maxima representing background signal or nuclear DNA were removed manually from the dataset. The resulting number of maxima represented the number of nucleoids in the given region of interest. Only budded cells with a b:m > 0.4 were scored, and cells with weak DAPI signal were discarded from the quantification of nucleoid numbers. The budded cells analyzed were used to determine the % cells with DAPI-marked mtDNA signal.

For post-cytokinetic mitochondrial volume analysis, cytokinetic events were manually identified on non-deconvolved images in FIJI. The same procedure as described above was performed for the first timepoint in which the Myo1-GFP signal was absent from the mother-bud neck.

Cell/mother/bud volume was determined from the radii of ellipse ROIs of mothers and buds (in µm) measured in FIJI. These data were transformed into an ellipsoid approximation of the ROI volume (in µm^3^) using the following equation: ellipsoid volume = (4/3)*π*(major radius/2)*(minor radius^2^/4).

### Plate Reader Growth Assay

Growth assays were performed in 96-well clear bottom plates (Nunclon Delta Surface – Thermo Scientific) with OD_600_ measurements taken every 10 minutes for 18.5 h at 30°C with no shaking on a SpectraMax iD3 (Molecular Devices) plate reader. Each plate was prepared with three technical replicates for each biological replicate (for which there are 3 per genotype) in YPD, and the plate was sealed with a clear breathable membrane to prevent evaporation. The change in OD_600_ per min. (slope) was calculated from a portion of the growth curve where WT displayed linear growth (between 150-270 min.). Outliers were determined using the following criteria: the endpoint OD_600_ was outside the upper and lower limits of the quartiles 1 and 3 (Q1 and Q3, respectively). The lower limit was calculated using Q1-[1.5*(Q3-Q1)]. The upper limit was calculated using Q3+[1.5*(Q3-Q1)].

### Statistical analyses

Data distributions were assumed to be normal. Either an unpaired two-tailed *t-test* or a one-way ordinary ANOVA with Dunnett’s multiple comparisons was performed as indicated in the figure legends. All statistical analyses, including linear regression, were performed using GraphPad Prism.

## ACKNOWLEDGEMENTS

We thank members of the Lackner lab for feedback on the manuscript and critical scientific discussions. We also thank Northwestern’s Cell Biology Supergroup and the Wignall-Lackner Cell Biology Group for constructive feedback on the project. Additional thanks to Jessica Hornick and Tong Zhang for assisting with the fluorescence microscopy. All microscopy was performed at the Biological Imaging Facility at Northwestern University (RRID:SCR_017767), supported by the Chemistry for Life Processes Institute, the Northwestern University Office for Research, the Department of Molecular Biosciences, and the Rice Foundation. Plasmids expressing the Myo2 point mutants were a kind gift from Lois Weisman at the University of Michigan.

M.W.R. was supported by the NIGMS Training Grant T32 GM008061 and the National Science Foundation Graduate Research Fellowship Program under Grant No. DGE-2234667. W.C. was supported by NIGMS Training Grant T32 GM008382. C.D. was supported in part by the Northwestern University Graduate School Cluster in Biotechnology, Systems, and Synthetic Biology, which is affiliated with the Biotechnology Training Program. E.M.R was supported by the NIGMS T32 GM008449 and the National Science Foundation Graduate Research Fellowship Program under Grant No. DGE-2234667. L.L.L. was supported by NIGMS grant R01 GM120303 and R56 AG088061. The findings and conclusions expressed in this material are those of the authors and do not necessarily reflect the views of the NIGMS or National Science Foundation.

---

**Figure S1.**
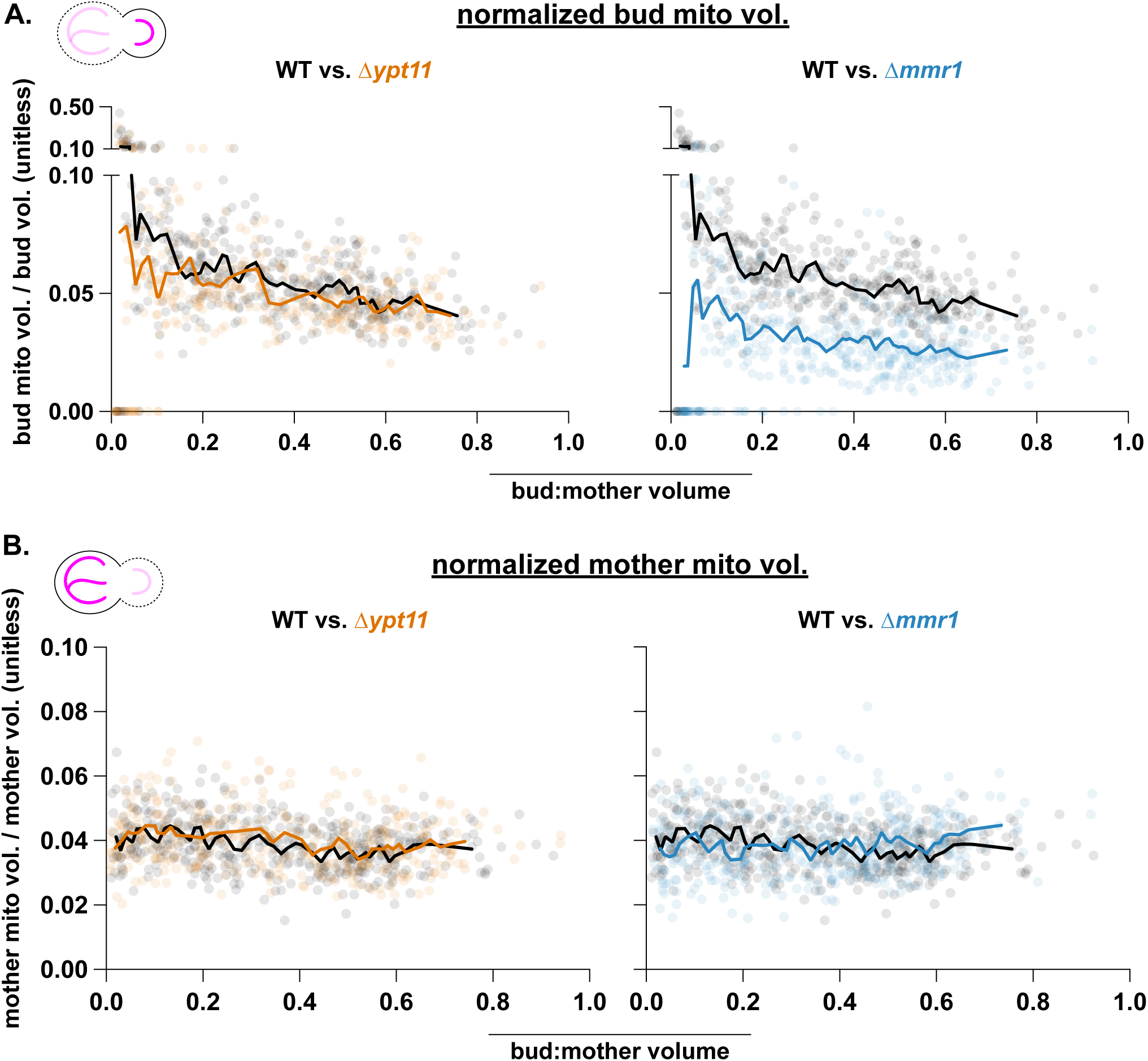
Normalized mitochondrial volumes of buds and mothers in mitochondrial inheritance adaptor mutants. The same dataset from Fig. 1 C was used to graph both the normalized mitochondrial volumes of buds (A) and mothers (B) over b:m volume. Data is represented as in Fig. 1C. The y-axes in (A) are split in two for data visualization purposes.

**Figure S2.**
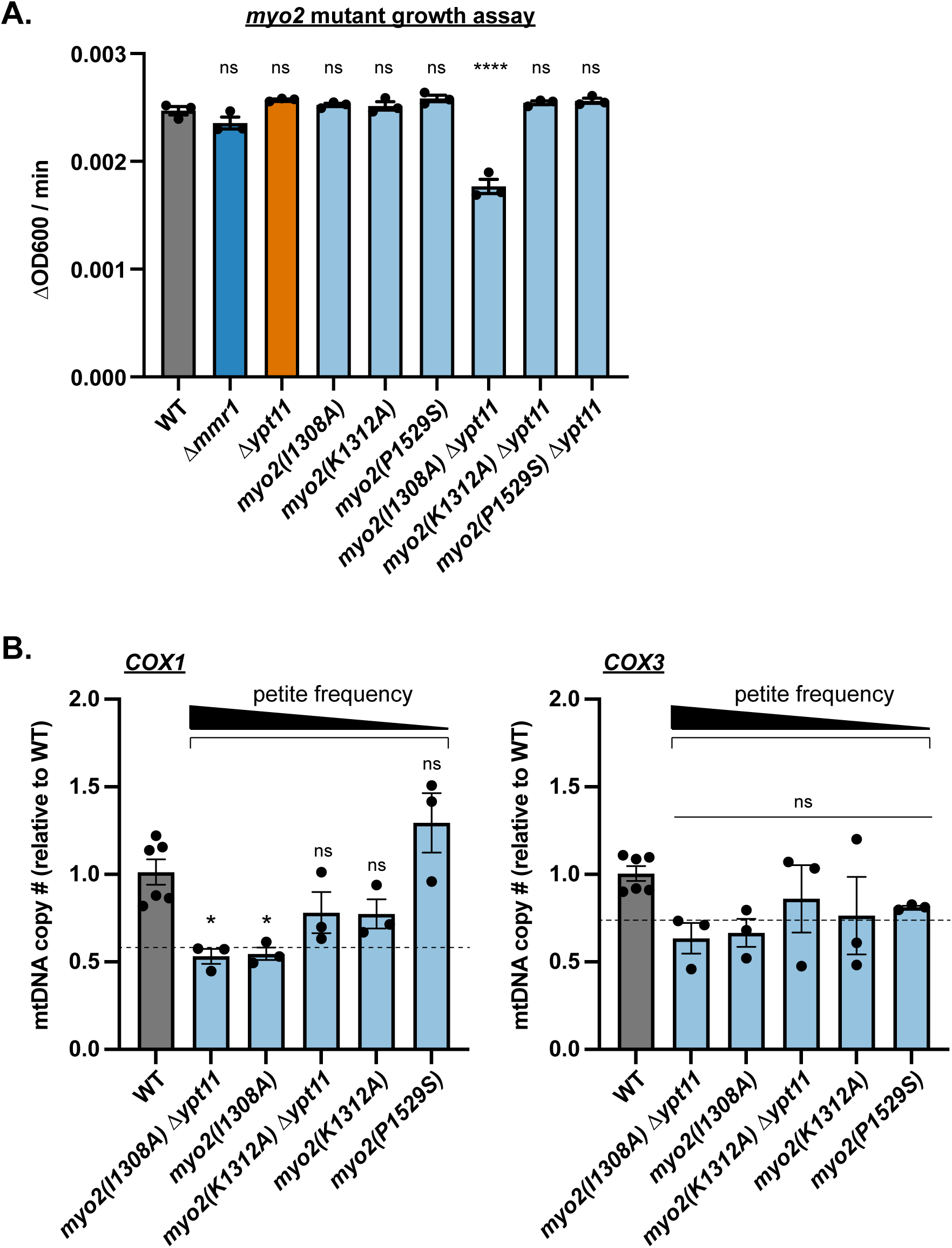
Characterization of strains with point mutations on Myo2-CBD that disrupt Mmr1 binding. (A) Growth assay of wild-type (WT), Δ*mmr1*, Δ*ypt11*, and *myo2* mutant strains. Cells were grown in fermentation conditions in a plate reader, and measurements of optical densities at 600 nm (OD600) were used to determine growth rate (ΔOD600/min). Each black dot represents 1 of 3 total biological replicates. Colored bars show grand mean with error bars representing ± SEM. p-values are in comparison to WT (* p < 0.05; ** p < 0.01; **** p < 0.0001; ordinary one-way ANOVA multiple comparisons). (B) qPCR analysis of *COX1* and *COX3* in *myo2* mutant strains was performed and is represented as in Fig. 3 E. Dashed black lines represent the grand mean of the Δ*mmr1* value (as determined in Fig. 3 E) for comparison. Mutants are shown in order of decreasing mean petite frequencies (from left to right) as determined in Fig. 4 D. p-values are in comparison to WT (* p < 0.05; ** p < 0.01; **** p < 0.0001; ordinary one-way ANOVA multiple comparisons).

**Supplemental Table 1.**

| LLY Number | Genotype | Source |
| --- | --- | --- |
| 92(A)/93(alpha) | W303 (*ade2-1; leu2-3; his3- 11,15; trp1-1; ura3-1; can1-100*) | Thomas and Rothstein, 1989 |
| 3584 | W303 *ADH::mito-DsRED::URA* | This study |
| 3585 | W303 Δ*mmr1::KAN ADH::mito-DsRED::URA* | This study |
| 3586 | W303 Δ*ypt11::KAN ADH::mito-DsRED::URA* | This study |
| 5408/5409 | W303 *ADH::mito-DsRED::URA MYO1-yEGFP::HIS* | This study |
| 5777 | W303 Δ*mmr1::KAN ADH::mito-DsRED::URA MYO1-yEGFP::HIS* | This study |
| 490 | W303 Δ*mmr1::HIS* | This study |
| 491/2852 | W303 Δ*mmr1::KAN* | Chen et al. 2018 (LLY491); this study (LLY2852) |
| 962/963 | W303 Δ*ypt11::NAT* | This study |
| 964/965 | W303 Δ*ypt11::KAN* | This study |
| 5012/5013 | W303 Δ*mrx6::HIS* | Wisniewski et al. 2024 |
| 6544/6545 | W303 *rho*<sup>0</sup> *ADH::mito-DsRED::URA* | This study |
| 6540/6541 | W303 *rho*<sup>0</sup> Δ*mmr1::KAN ADH::mito-DsRED::URA* | This study |
| 6538/6539 | W303 *rho*<sup>0</sup> Δ*ypt11::NAT ADH::mito-DsRED::URA* | This study |
| 1543 | W303 *mmr1*Δ*mito-BD [MMR1(Δ76-195)-yEGFP::HIS]* | Chen et al. 2018 |
| 2618 | W303 *mmr1*<sup>4E</sup> [*mmr1*(R80E/R86E/K95E/K98E)-yEGFP::HIS] | Chen et al. 2018 |
| 4043/4044 | W303 *myo2*(P1529S)::HIS | This study |
| 4136/4137 | W303 *myo2*(I1308A)::HIS | This study |
| 4260 | W303 *myo2*(P1529S)::HIS Δ*ypt11::NAT* | This study |
| 4261 | W303 *myo2*(I1308A)::HIS Δ*ypt11::NAT* | This study |
| 4262/4263 | W303 *myo2*(K1312A)::HIS | This study |
| 4284 | W303 *myo2*(K1312A)::HIS Δ*ypt11::NAT* | This study |
| 4751/4753 | W303 *myo2*(I1308A)::HIS *ADH::mito-DsRED::URA* | This study |
| 5971/5972 | W303 *myo2*(K1312A)::HIS Δ*ypt11::NAT ADH::mito-DsRED::URA* | This study |
| 5973/5974/5975 | W303 *myo2*(I1308A)::HIS Δ*ypt11::NAT ADH::mito-DsRED::URA* | This study |
| 5976/5977/5978 | W303 *myo2*(K1312A)::HIS *ADH::mito-DsRED::URA* | This study |
| 6680/6681 | W303 *myo2*(P1529S)::HIS *ADH::mito-DsRED::URA* | This study |
| 5476/5477 | W303 Δ*mmr1::KAN* Δ*sml1::HIS* | This study |
| 6530/6531/6683 | W303 Δ*mmr1::KAN* Δ*sml1::HIS ADH::mito-DsRED::URA* | This study |
| 5358/5359 | W303 Δ*mmr1::KAN* Δ*mrx6::HIS* | This study |
| 6542/6682 | W303 Δ*mmr1::KAN* Δ*mrx6::HIS ADH::mito-DsRED::URA* | This study |
| 5479/5480/5481 | W303 Δ*mmr1::KAN* Δ*cim1::HIS* | This study |
| 6534/6535/6536/6537 | W303 $\Delta mmr1::KAN$ $\Delta cim1::HIS$ $ADH::mito-DsRED::URA$ | This study |

**Supplemental Table 2.**
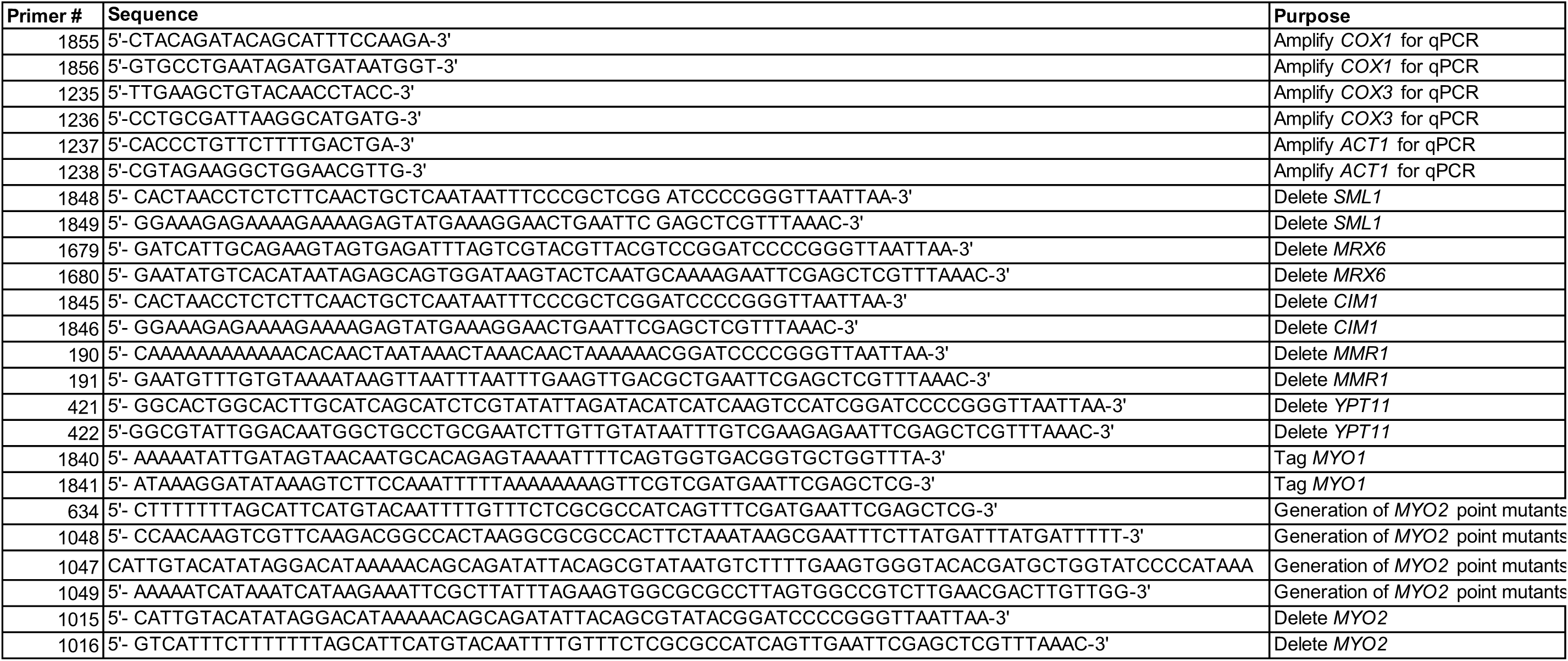

**Supplemental Table 3.**

| **LLEC#** | **Plasmid Name** | **Source** |
| --- | --- | --- |
| 776 | pRS306 | Sikorski and Hieter, 1989 |
| 806 | pRS306 *ADH::mito-DsRED::URA* | This study |
| 27 | pFA6a-kanMX6 | Longtine et al. 1998 |
| 54 | pKT128 pFA6a-link-yEGFP-SpHIS5 | Sheff and Thorn 2004 |
| 29 | pFA6a-His3MX6 | Longtine et al. 1998 |
| 473 | pFA6a-NAT | Goldstein and McCusker 1999 |
| 668 | pRS413 *myo2(K1312A)* | Eves et al. 2012 |
| 669 | pRS413 *myo2(P1529S)* | Eves et al. 2012 |
| 670 | pRS413 *myo2(I1308A)* | Eves et al. 2012 |

## REFERENCES

Anderson, A.P., X. Luo, W. Russell, and Y.W. Yin. 2020. Oxidative damage diminishes mitochondrial DNA polymerase replication fidelity. Nucleic Acids Res. 48:817–829.

Arai, S., Y. Noda, S. Kainuma, I. Wada, and K. Yoda. 2008. Ypt11 functions in bud-directed transport of the Golgi by linking Myo2 to the coatomer subunit Ret2. Curr Biol. 18:987–991.

Bockler, S., X. Chelius, N. Hock, T. Klecker, M. Wolter, M. Weiss, R.J. Braun, and B. Westermann. 2017. Fusion, fission, and transport control asymmetric inheritance of mitochondria and protein aggregates. J Cell Biol. 216:2481–2498.

Boldogh, I.R., S.L. Ramcharan, H.C. Yang, and L.A. Pon. 2004. A type V myosin (Myo2p) and a Rab-like G-protein (Ypt11p) are required for retention of newly inherited mitochondria in yeast cells during cell division. Mol Biol Cell. 15:3994–4002.

Chen, W., H.A. Ping, and L.L. Lackner. 2018. Direct membrane binding and self-interaction contribute to Mmr1 function in mitochondrial inheritance. Mol Biol Cell. 29:2346–2357.

Chernyakov, I., F. Santiago-Tirado, and A. Bretscher. 2013. Active segregation of yeast mitochondria by Myo2 is essential and mediated by Mmr1 and Ypt11. Curr Biol. 23:1818–1824.

Eves, P.T., Y. Jin, M. Brunner, and L.S. Weisman. 2012. Overlap of cargo binding sites on myosin V coordinates the inheritance of diverse cargoes. J Cell Biol. 198:69–85.

Filograna, R., C. Koolmeister, M. Upadhyay, A. Pajak, P. Clemente, R. Wibom, M.L. Simard, A. Wredenberg, C. Freyer, J.B. Stewart, and N.G. Larsson. 2019. Modulation of mtDNA copy number ameliorates the pathological consequences of a heteroplasmic mtDNA mutation in the mouse. Sci Adv. 5:eaav9824.

Filograna, R., M. Mennuni, D. Alsina, and N.G. Larsson. 2021. Mitochondrial DNA copy number in human disease: the more the better? FEBS Lett. 595:976–1002.

Fox, T.D., L.S. Folley, J.J. Mulero, T.W. McMullin, P.E. Thorsness, L.O. Hedin, and M.C. Costanzo. 1991. Analysis and manipulation of yeast mitochondrial genes. Methods Enzymol. 194:149–165.

Friedman, J.R., and J. Nunnari. 2014. Mitochondrial form and function. Nature. 505:335–343.

Goke, A., S. Schrott, A. Mizrak, V. Belyy, C. Osman, and P. Walter. 2020. Mrx6 regulates mitochondrial DNA copy number in Saccharomyces cerevisiae by engaging the evolutionarily conserved Lon protease Pim1. Mol Biol Cell. 31:527–545.

Goldstein, A.L., and J.H. McCusker. 1999. Three new dominant drug resistance cassettes for gene disruption in Saccharomyces cerevisiae. Yeast. 15:1541–1553.

Higuchi-Sanabria, R., J.K. Charalel, M.P. Viana, E.J. Garcia, C.N. Sing, A. Koenigsberg, T.C. Swayne, J.D. Vevea, I.R. Boldogh, S.M. Rafelski, and L.A. Pon. 2016. Mitochondrial anchorage and fusion contribute to mitochondrial inheritance and quality control in the budding yeast Saccharomyces cerevisiae. Mol Biol Cell. 27:776–787.

Itoh, T., E.A. Toh, and Y. Matsui. 2004. Mmr1p is a mitochondrial factor for Myo2p-dependent inheritance of mitochondria in the budding yeast. EMBO J. 23:2520–2530.

Itoh, T., A. Watabe, E.A. Toh, and Y. Matsui. 2002. Complex formation with Ypt11p, a rab-type small GTPase, is essential to facilitate the function of Myo2p, a class V myosin, in mitochondrial distribution in Saccharomyces cerevisiae. Mol Cell Biol. 22:7744–7757.

Jajoo, R., Y. Jung, D. Huh, M.P. Viana, S.M. Rafelski, M. Springer, and J. Paulsson. 2016. Accurate concentration control of mitochondria and nucleoids. Science. 351:169–172.

Janke, C., M.M. Magiera, N. Rathfelder, C. Taxis, S. Reber, H. Maekawa, A. Moreno-Borchart, G. Doenges, E. Schwob, E. Schiebel, and M. Knop. 2004. A versatile toolbox for PCR-based tagging of yeast genes: new fluorescent proteins, more markers and promoter substitution cassettes. Yeast. 21:947–962.

Jiang, M., T.E.S. Kauppila, E. Motori, X. Li, I. Atanassov, K. Folz-Donahue, N.A. Bonekamp, S. Albarran-Gutierrez, J.B. Stewart, and N.G. Larsson. 2017. Increased Total mtDNA Copy Number Cures Male Infertility Despite Unaltered mtDNA Mutation Load. Cell Metab. 26:429–436 e424.

Kraft, L.M., and L.L. Lackner. 2017. Mitochondria-driven assembly of a cortical anchor for mitochondria and dynein. J Cell Biol. 216:3061–3071.

Lackner, L.L. 2014. Shaping the dynamic mitochondrial network. BMC Biol. 12:35.

Lackner, L.L., H. Ping, M. Graef, A. Murley, and J. Nunnari. 2013. Endoplasmic reticulum-associated mitochondria-cortex tether functions in the distribution and inheritance of mitochondria. Proc Natl Acad Sci U S A. 110:E458–467.

Lewandowska, A., J. Macfarlane, and J.M. Shaw. 2013. Mitochondrial association, protein phosphorylation, and degradation regulate the availability of the active Rab GTPase Ypt11 for mitochondrial inheritance. Mol Biol Cell. 24:1185–1195.

Li, K.W., M.S. Lu, Y. Iwamoto, D.G. Drubin, and R.T.A. Pedersen. 2021. A preferred sequence for organelle inheritance during polarized cell growth. J Cell Sci. 134:jcs258856.

Lippincott, J., and R. Li. 1998. Sequential assembly of myosin II, an IQGAP-like protein, and filamentous actin to a ring structure involved in budding yeast cytokinesis. J Cell Biol. 140:355–366.

Livak, K.J., and T.D. Schmittgen. 2001. Analysis of relative gene expression data using real-time quantitative PCR and the 2(-Delta Delta C(T)) Method. Methods. 25:402–408.

Longtine, M.S., A. McKenzie, 3rd, D.J. Demarini, N.G. Shah, A. Wach, A. Brachat, P. Philippsen, and J.R. Pringle. 1998. Additional modules for versatile and economical PCR-based gene deletion and modification in Saccharomyces cerevisiae. Yeast. 14:953–961.

Miettinen, T.P., and M. Bjorklund. 2016. Cellular Allometry of Mitochondrial Functionality Establishes the Optimal Cell Size. Dev Cell. 39:370–382.

Miyakawa, I. 2017. Organization and dynamics of yeast mitochondrial nucleoids. Proc Jpn Acad Ser B Phys Biol Sci. 93:339–359.

Nishiyama, S., H. Shitara, K. Nakada, T. Ono, A. Sato, H. Suzuki, T. Ogawa, H. Masaki, J. Hayashi, and H. Yonekawa. 2010. Over-expression of Tfam improves the mitochondrial disease phenotypes in a mouse model system. Biochem Biophys Res Commun. 401:26–31.

Ogur, M., and R. St John. 1956. A differential and diagnostic plating method for population studies of respiration deficiency in yeast. J Bacteriol. 72:500–504.

Osman, C., T.R. Noriega, V. Okreglak, J.C. Fung, and P. Walter. 2015. Integrity of the yeast mitochondrial genome, but not its distribution and inheritance, relies on mitochondrial fission and fusion. Proc Natl Acad Sci U S A. 112:E947–956.

Pekkurnaz, G., and X. Wang. 2022. Mitochondrial heterogeneity and homeostasis through the lens of a neuron. Nat Metab. 4:802–812.

Rafelski, S.M., M.P. Viana, Y. Zhang, Y.H. Chan, K.S. Thorn, P. Yam, J.C. Fung, H. Li, F. Costa Lda, and W.F. Marshall. 2012. Mitochondrial network size scaling in budding yeast. Science. 338:822–824.

Roussou, R., D. Metzler, F. Padovani, F. Thoma, R. Schwarz, B. Shraiman, K.M. Schmoller, and C. Osman. 2024. Real-time assessment of mitochondrial DNA heteroplasmy dynamics at the single-cell level. EMBO J. 43:5340–5359.

Schindelin, J., I. Arganda-Carreras, E. Frise, V. Kaynig, M. Longair, T. Pietzsch, S. Preibisch, C. Rueden, S. Saalfeld, B. Schmid, J.Y. Tinevez, D.J. White, V. Hartenstein, K. Eliceiri, P. Tomancak, and A. Cardona. 2012. Fiji: an open-source platform for biological-image analysis. Nat Methods. 9:676–682.

Schrott, S., and C. Osman. 2023. Two mitochondrial HMG-box proteins, Cim1 and Abf2, antagonistically regulate mtDNA copy number in Saccharomyces cerevisiae. Nucleic Acids Res. 51:11813–11835.

Seel, A., F. Padovani, M. Mayer, A. Finster, D. Bureik, F. Thoma, C. Osman, T. Klecker, and K.M. Schmoller. 2023. Regulation with cell size ensures mitochondrial DNA homeostasis during cell growth. Nat Struct Mol Biol. 30:1549–1560.

Shadel, G.S. 1999. Yeast as a model for human mtDNA replication. Am J Hum Genet. 65:1230–1237.

Sheff, M.A., and K.S. Thorn. 2004. Optimized cassettes for fluorescent protein tagging in Saccharomyces cerevisiae. Yeast. 21:661–670.

Sikorski, R.S., and P. Hieter. 1989. A system of shuttle vectors and yeast host strains designed for efficient manipulation of DNA in Saccharomyces cerevisiae. Genetics. 122:19–27.

Smith, K.K., J.D. Moreira, C.R. Wilson, J.O. Padera, A.N. Lamason, L. Xue, D.M. Gopal, D.B. Flynn, and J.L. Fetterman. 2024. A systematic review on the biochemical threshold of mitochondrial genetic variants. Genome Res. 34:341–365.

Swayne, T.C., C. Zhou, I.R. Boldogh, J.K. Charalel, J.R. McFaline-Figueroa, S. Thoms, C. Yang, G. Leung, J. McInnes, R. Erdmann, and L.A. Pon. 2011. Role for cER and Mmr1p in anchorage of mitochondria at sites of polarized surface growth in budding yeast. Curr Biol. 21:1994–1999.

Tang, K., Y. Li, C. Yu, and Z. Wei. 2019. Structural mechanism for versatile cargo recognition by the yeast class V myosin Myo2. J Biol Chem. 294:5896–5906.

Taylor, S.D., H. Zhang, J.S. Eaton, M.S. Rodeheffer, M.A. Lebedeva, W. O’Rourke T, W. Siede, and G.S. Shadel. 2005. The conserved Mec1/Rad53 nuclear checkpoint pathway regulates mitochondrial DNA copy number in Saccharomyces cerevisiae. Mol Biol Cell. 16:3010–3018.

Thomas, B.J., and R. Rothstein. 1989. Elevated recombination rates in transcriptionally active DNA. Cell. 56:619–630.

Viana, M.P., S. Lim, and S.M. Rafelski. 2015. Quantifying mitochondrial content in living cells. Methods Cell Biol. 125:77–93.

Wisniewski, B.T., J.C. Casler, and L.L. Lackner. 2024. Significantly reduced, but balanced, rates of mitochondrial fission and fusion are sufficient to maintain the integrity of yeast mitochondrial DNA. Mol Biol Cell. 35:br25.

Zwonitzer, K.D., L.G. Tressel, Z. Wu, S. Kan, A.K. Broz, J.P. Mower, T.A. Ruhlman, R.K. Jansen, D.B. Sloan, and J.C. Havird. 2024. Genome copy number predicts extreme evolutionary rate variation in plant mitochondrial DNA. Proc Natl Acad Sci U S A. 121:e2317240121.

